# Sequencing, fast and slow: profiling microbiomes in human samples with nanopore sequencing

**DOI:** 10.1101/2023.05.18.541257

**Authors:** Yunseol Park, Jeesu Lee, Hyunjin Shim

**Author notes:** These authors have contributed equally to this work (Yunseol Park, Jeesu Lee). Corresponding Author: Hyunjin Shim. Contributions Experiments were primarily conducted by H.S. Analyses were primarily conducted by Y.P., J.L., and H.S. Specifically, EPI2ME analyses were performed by H.S., classified nanopore analyses were led by Y.P., and unclassified nanopore analyses were conducted by J.L. and H.S. The study was conceived by H.S., and all authors contributed to writing the manuscript.

## Abstract

Rapid and accurate pathogen identification is crucial in effectively combating infectious diseases. However, the current diagnostic tools for bacterial infections predominantly rely on century-old culture-based methods. Furthermore, recent research highlights the significance of host-microbe interactions within the host microbiota in influencing the outcome of infection episodes. As our understanding of science and medicine continues to advance, there is a pressing need for innovative diagnostic methods that can identify pathogens and also rapidly and accurately profile the microbiome landscape in human samples. In clinical settings, such diagnostic tools will become a powerful predictive instrument in directing the diagnosis and prognosis of infectious diseases by providing comprehensive insights into the patient’s microbiota. Here, we explore the potential of long-read sequencing in profiling the microbiome landscape from various human samples in terms of speed and accuracy. Using nanopore sequencers, we generate native DNA sequences from saliva and stool samples rapidly, from which each long-read is basecalled in real-time to provide downstream analyses such as taxonomic classification and antimicrobial resistance through the built-in software (< 12 hours). Subsequently, we utilize the nanopore sequence data for in-depth analysis of each microbial species in terms of host-microbe interaction types and deep learning-based classification of unidentified reads. We find that the nanopore sequence data encompass complex information regarding the microbiome composition of the host and its microbial communities, and also shed light on the unexplored human mobilome including bacteriophages. In this study, we use two different systems of long-read sequencing to give insights into human microbiome samples in the ‘slow’ and ‘fast’ modes, which raises additional inquiries regarding the precision of this novel technology and the feasibility of extracting native DNA sequences from other human microbiomes.

## Introduction

Rapid and accurate identification of pathogens is crucial for effectively treating and managing infectious diseases. Traditional diagnostic methods for bacterial infections, such as culture-based techniques, have been largely unchanged in clinical practice for several decades [1,2]. However, these methods often take several days for identification and susceptibility testing of bacterial pathogens and are prone to false-negative results during antimicrobial therapy. For example, a recent study showed that the median time to pathogen identification for bloodstream infections using traditional culture-based methods takes around three days [3]. This delay in diagnostic procedures can result in inappropriate antibiotic therapy, which can negatively affect patient outcomes and lead to antibiotic resistance development [1]. Furthermore, culture-based techniques may not detect all bacterial infections, particularly if the patient is undergoing antimicrobial therapy.

Recent evidence suggests that it is important to have a comprehensive view of the microbial communities in which the pathogen resides to predict the progress of infection in clinical settings [4]. The human microbiota, consisting of a vast array of microorganisms, including bacteria, viruses, fungi, and protozoa, colonizes many different niches within the human body. Host-microbe interactions within the host microbiota play a vital role in determining the growth and establishment of pathogenic microbes. These interactions can range from beneficial to commensal to pathogenic, and a subtle shift in the balance of these interactions can have profound effects on the host’s health, including inflammatory bowel disease [5] and neurological disorders [6]. Recent research has highlighted the role of the host microbiota in shaping the host’s immune response to pathogenic microorganisms, both through direct interactions with the immune system and through modulation of the host’s innate and adaptive immune responses [7]. Studies have demonstrated that the host microbiota can provide colonization resistance against invading pathogens, limiting their growth and preventing their establishment within the host [8]. Furthermore, the host microbiota can also impact the virulence of pathogenic microorganisms through a variety of mechanisms, including competition for nutrients, secretion of antimicrobial compounds, and modulation of the expression of virulence factors [9]. Alterations in the composition of the host microbiota, such as those caused by antibiotics or changes in diet, can disrupt these finely balanced host-microbe interactions, leading to increased susceptibility to infections [10].

Understanding the intricate interactions between the host microbiota and pathogenic microorganisms is critical for the development of effective treatments for infectious diseases. This knowledge can inform the development of novel therapeutics that target specific bacterial species or modulate the host’s immune response to promote the restoration of healthy microbiota and reduce the risk of disease [11]. Alterations in the composition of the microbiota can disrupt these interactions, leading to increased susceptibility to infections. Currently, we are in need of novel diagnostic methods that can both identify pathogens and profile the microbiome landscape in human samples rapidly and accurately. Given the expanding knowledge of the role of the microbiome in human health, this feature is a substantial advancement to traditional diagnostic methods that focus primarily on pathogen identification, such as culture-based diagnosis and MALDI-TOF mass spectrometry fingerprinting [1].

Molecular-based diagnostic methods, such as polymerase chain reaction (PCR) and next-generation sequencing (NGS), can detect a wider range of bacterial pathogens with greater sensitivity and specificity [12]. In recent years, the development of such technologies has enabled the rapid and accurate identification of pathogens. Particularly, NGS can generate vast amounts of sequence data in a short amount of time, providing rapid and accurate results for pathogen identification that enables clinicians to make informed decisions about antimicrobial treatments [13]. Several studies have shown the potential of NGS in clinical settings for the rapid identification of bacterial infections, such as the rapid identification of a bacterial outbreak in a neonatal intensive care unit [14] and in patients with sepsis [15]. Despite the potential of NGS, the technology is still not widely available in clinical settings, and there are several challenges related to quality control measures that need to be overcome. These challenges include the need for standardized protocols for sample preparation, sequencing, and data analysis, particularly since these methods produce short-read DNA sequences in metagenomic samples that require elaborate bioinformatic reconstructions [16].

Most recently, long-read sequencing technologies are revolutionizing genomics research by producing high-throughput sequencing of DNA reads longer than those obtained by traditional short-read sequencing methods. Nanopore sequencing is one of the long-read sequencing technologies, in which a DNA molecule is passed through a nanopore, and the electrical signal generated by the movement of the nucleotides through the pore is used to determine the sequence of the DNA molecule [17]. The development of long-read sequencing has had a significant impact on genomics research and has enabled the study of complex genomes, structural variation, and epigenetic modifications with unprecedented accuracy and resolution [18,19]. Long-read sequencing has the potential to further enhance the field of metagenomics by enabling the identification and characterization of unculturable microbes and the study of host-microbe interactions in complex microbial communities [20]. A recent study demonstrated the potential of nanopore sequencing as the clinical diagnosis of bacterial lower respiratory infections by directly sequencing sputum samples, providing comparable results to culture-based methods, but with significantly faster turnaround times [21,22].

In this study, we investigate the potential of long-read sequencing as a futuristic diagnostic tool to rapidly profile the microbiome landscape of diverse human samples. We use two different modes to sequence long-read native DNA sequences from diverse human microbiomes, including saliva and stool samples. The first part consists of a ‘fast’ mode, which aims to generate a biological interpretation of nanopore sequencing within 12 hours from sampling to data analysis (Figure 1). This fast mode is aimed at providing ultrarapid analysis of nanopore sequence data such as pathogen identification and antimicrobial resistance (AMR) under a clinical scenario of tight time constraints where the streamlined pipeline of a diagnostic tool is essential for effective treatment. This mode is fast and automatic, and it enables clinicians to make quick identifications and decisions on antibiotic therapy based on the pathogen and its related microbes without much investment of time and effort. A ‘slow’ mode is aimed at providing deeper insight into the microbiome landscape of a patient for prognostic purposes, during which the microbial communities are analyzed more rigorously. This mode is slow and deliberate, and it engages the researchers to predict the long-term trajectory of an infection outcome using complex information such as host-microbe interactions and deep learning-based classification of unknown organisms. Overall, we aim to provide a comprehensive and insightful view of long-read sequencing as an innovative diagnostic tool for bacterial infections by rapidly profiling the microbiome landscape from various human samples.

**Fig. 1:**
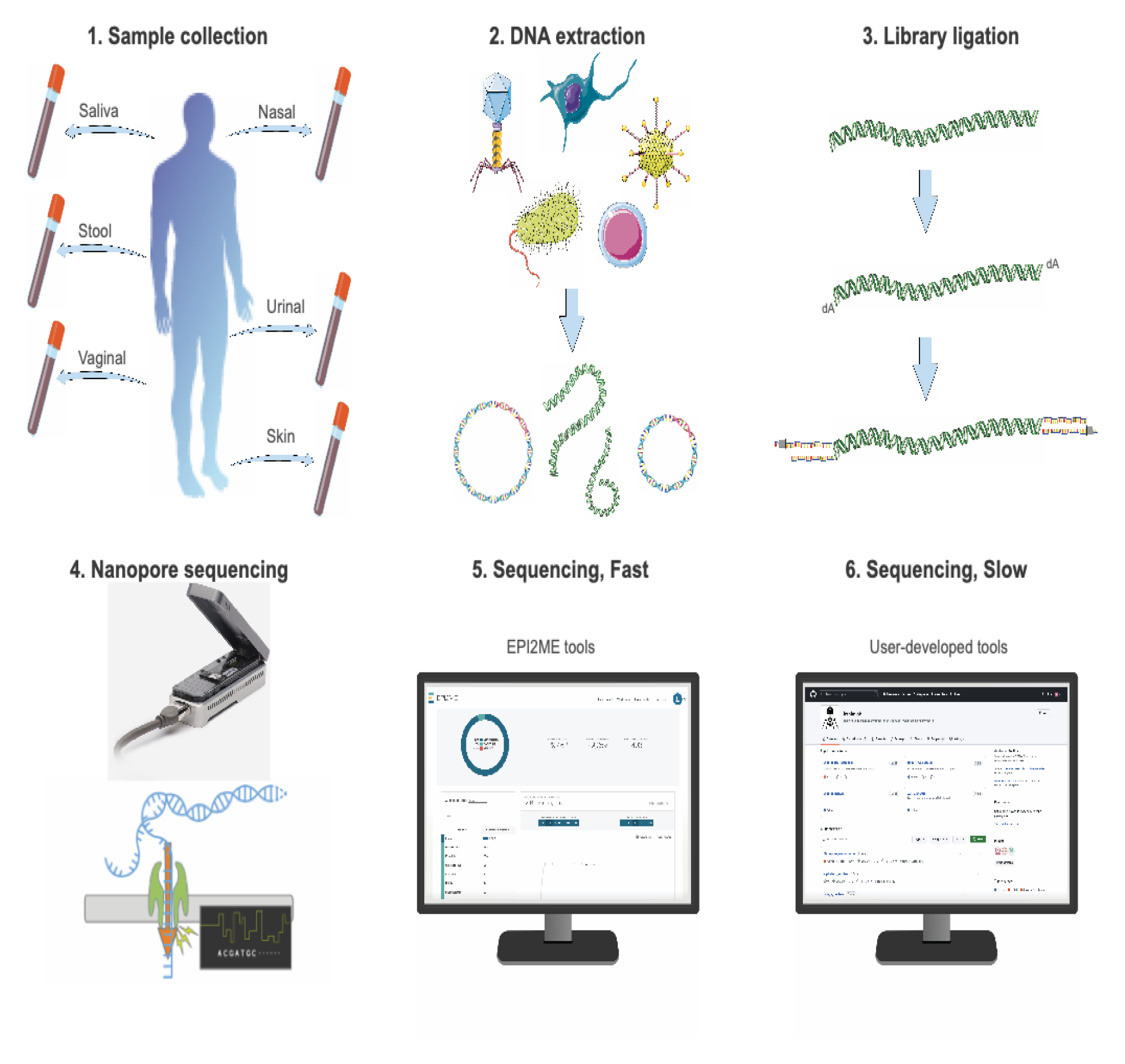
Experimental setup and process of nanopore sequencing. The sample collection of human microbiome is collected with a sterile specimen swab. Native DNA of the metagenomic sample is extracted with a commercial kit with minimum shearing to obtain high molecular weight DNA. The extracted DNA is ligated with a library kit provided by Oxford Nanopore that has been optimized for sequencing native and long-read DNAs in Flongle flow cells. Flongle flow cells are fitted with an adaptor to a nanopore sequencer for rapid and cost-effective tests, running for at least 12 hours to maximize yield per sample. Basecalling is done in real-time for a fast sequencing mode, during which results can be obtained using cloud-based software tools for taxonomic classification and AMR analysis. For a slow sequencing mode, user-developed tools can be used to conduct exploratory analysis on the same nanopore data, including host-microbe interaction assessment and deep learning-based classification of unidentified reads.

## Results

### Native DNA could be extracted and sequenced from the saliva and stool samples

We used the QIAamp DNA Microbiome Kit to extract the native DNA of various human samples from healthy volunteers (Figure 1). The minimum DNA quantity needed for native DNA direct sequencing using Flongle flow cells is 500 ng according to the manufacturer’s protocol, and the lowest DNA concentration achieved for the saliva and stool samples was 20.5 ng/µl, yielding sufficient quantities for nanopore sequencing (Table S1). However, the same kit was found to be ineffective for extracting native DNA from urine, nasal, and vaginal samples sourced from healthy volunteers, failing to provide the minimum DNA quantities required for nanopore sequencing (Table S1). These findings suggest that the QIAamp DNA Microbiome Kit may have limited utility for extracting native DNA from certain human microbiome sources, and alternative DNA extraction methods may need to be explored for these sample types.

Another disadvantage of using the QIAamp DNA Microbiome Kit to extract native DNA from various microbiome samples comes from the fragmentation of DNAs during the extraction step (Figure 1). Fragmentation of DNA during extraction can be caused by a variety of factors, including mechanical and enzymatic shearing. The QIAamp DNA Microbiome Kit uses a bead-beating step to lyse cells, which can result in excessive mechanical shearing of DNA. Due to the DNA fragmentation, the shortest and the longest estimated N50 values are 378 bases and 1,090 bases, respectively (Table S2, Figure S1). N50 is a statistical measure commonly used in DNA sequencing to describe the quality of an assembly, and in the context of long-read sequencing, it is defined as the length of the shortest read within the set of the longest reads that constitute at least 50% of the sample [23]. Previously, it was shown that nanopore sequencers can produce long reads of around 10–30 kilobases (kb) reads in a typical sequencing experiment, while ultralong reads were shown to be around 3 megabases (Mb) with the N50 value of more than 100 kb [24].

### Fast sequencing shows oral and gut microbiomes have diverse microbial species

In this study, we ran one Flongle flow cell with each sample replicate for at least 12 hours, to exhaust the capacity of nanopores to obtain as many DNA reads in each run as possible (Figure S2). Depending on the sample, nanopore sequencing using Flongle flow cells was saturated as early as 3 hours (Figure S2; Saliva3_R2) and as late as 20 hours (Figure S2; Stool1_R2). The recommended hours of sequencing for nanopore Flongle can vary depending on the desired experimental output. For example, a recent study reported using Flongle flow cells with a sequencing time of 24 hours to generate high-quality, near-complete bacterial genomes of *Mycoplasma bovis* [25]. Similarly, another study utilized Flongle flow cells in a similar timeline to achieve high-quality, near-complete SARS-CoV-2 genome assemblies [26].

In this study, we used the high-accuracy basecalling program integrated into the MinKNOW software in real-time, with a minimum Q-score of 9. The Phred score and Quality score (Q-score) are both measures of the quality of sequencing data, where the Phred score is a logarithmic measure of the error probability originated to identify fluorescently labeled DNA bases by comparing observed and expected chromatogram peak shapes and resolution [27], widely used in Sanger sequencing and Illumina sequencing. For nanopore sequencing, per-nucleotide quality scores are based on the outputs of the neural networks used to produce the basecall. Q-scores are per-read quality scores, calculated by averaging the per-nucleotide quality scores and by expressing on the Phred scale [28]. Importantly, Q-scores consider that the error rate in nanopore sequencing is not constant across the read, and can vary depending on factors such as the sequence context and the quality of the signal.

After a complete sequencing run, we used the basecalled output for the quantitative and real-time identification of microbiome species from these metagenomic samples using the cloud-based data analysis platform (Table S3). This data analysis platform leverages long-read sequencing to enable the comparison of each read against databases containing reference genomes of bacteria, archaea, viruses, and fungi, achieved by constructing an indexing scheme that facilitates efficient searches of sequenced reads [29]. It rapidly classified and identified diverse species in each microbiome sample, even to the resolution of different strains of bacterial species (Table S3). The data analysis platform also rapidly determined the most reliable placement of these organisms in the taxonomy tree, assigning a score to each taxonomic placement (Figure S3).

The gut microbiomes contained the most number of species, while the oral microbiome contained varying amounts of microbial species (Table S3). For example, the Stool1 and Stool2 samples had more than 1,000 and 500 known microbial species present, respectively. The most abundant species consists of *Lactobacillus ruminis* in Stool1 and *Megamonas funiformis* in Stool2. Among these abundant species, it was notable that the gut microbiome from Stool1 contained most bacteria species widely known to be beneficial [30], whereas that from Stool2 had most bacterial species recently found to be commensal [31]. For the saliva samples, both the diversity and number of microbial species were lower and the role of each species in the host-microbe interaction was less obvious (Table S3). The most abundant bacterial species include *Haemophilus parainfluenzae* in Saliva1 and Saliva2, whereas *Rothia mucilaginosa* in Saliva 3. It was notable that the saliva microbiome from Saliva1 and Saliva2 contained most bacteria species widely known to be beneficial [32], whereas that from Saliva3 had most bacterial species recently found to be harmful [33]. Another notable observation includes the presence of viruses in these microbiome samples despite the use of an extraction kit that was not optimized for viral DNA extraction. The DNA reads that were classified as bacteriophage were of particular interest as the role and impact of these biological entities are just starting to get noticed in microbiome studies [34,35]. The most abundant virus species include Faecalibacterium phage in Stool1, crAss-like phage [36] in Stool2, Streptococcus phage in Saliva1 and Saliva2, and Shigella phage in Saliva 3 (Table S3). These first analyses show the diversity and abundance of microbial communities in human samples could be rapidly profiled - however, we conducted more in-depth analyzes of the host-microbe interaction types subsequently (see below).

In the saliva and stool samples, varying amounts of human DNAs were present despite the host DNA depletion step of the QIAamp DNA Microbiome Kit. In the stool samples, the microbiome DNA was enriched compared to the human DNA, with almost 100% of reads classified as bacterial species in Stool1 (Table 1). However, the saliva samples tend to have a lower percentage of the microbiome DNA in the sequencing output, with almost 90% of reads classified as eukaryotic species in Saliva1 and Saliva2 (Table 1). The third saliva sample diluted with 1000 µl of PBS solution had a better percentage content of the microbiome DNA, which shows that the quantity of a sample does not always correlate with the quality of reads in long-read sequencing. We studied these human DNA reads to assess if they could provide valuable information about the host, such as some genetic markers that could give alternative insight into the host-microbe interactions, but the yield output of a Flongle flow cell with a maximum 2.8 Gb was not enough to generate any significant coverage (unpublished). However, more high-throughput flow cells such as MinION and PromethION with a maximum output of 50 Gb and 290 Gb per flow cell, respectively, may be utilized to generate genomic data of the host and microbiome simultaneously, which may provide the most comprehensive view of the host-microbe interactions, given the recent findings of the interdependence of microbiome genomes and human genomes [37,38].

**Table 1.**
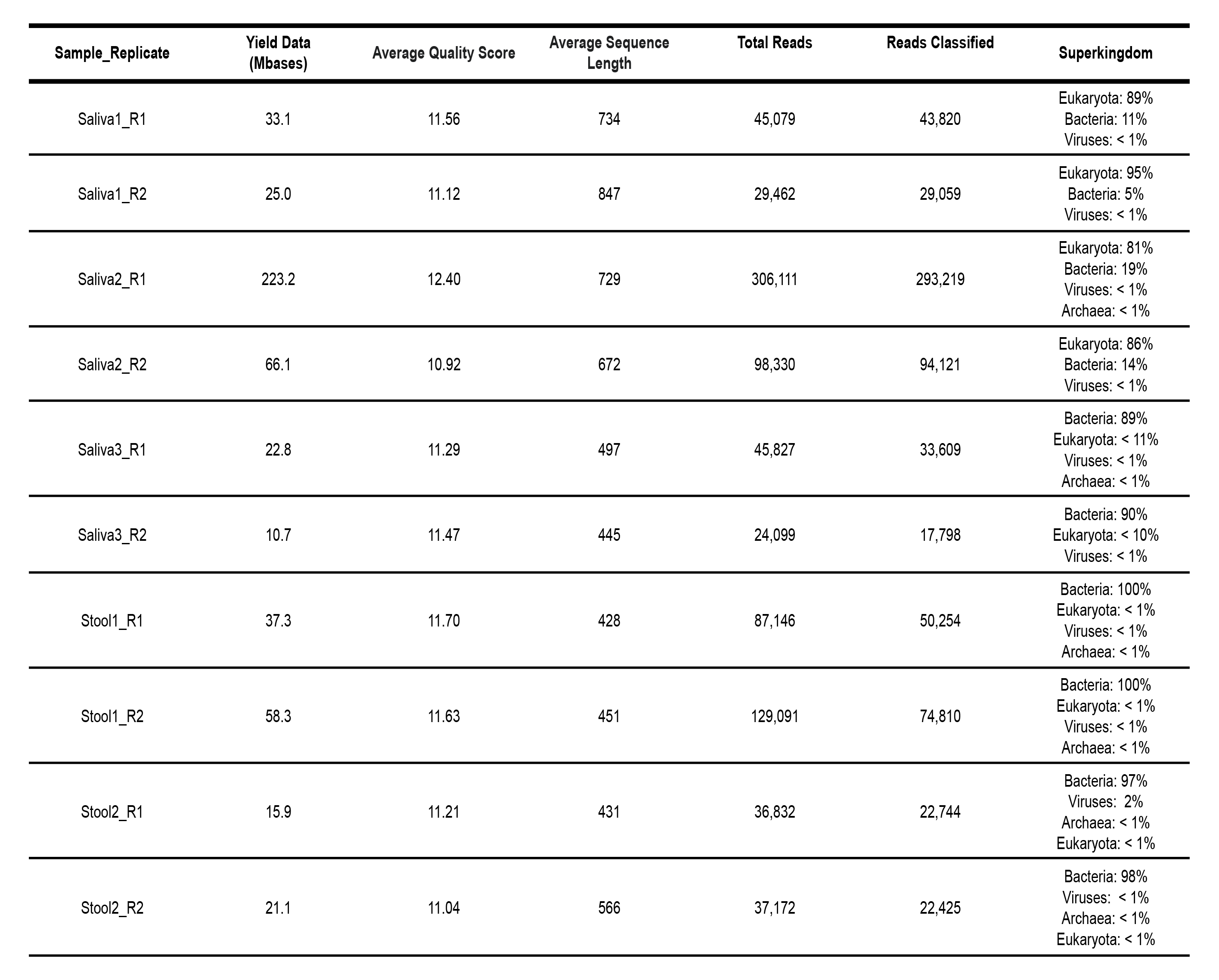
The real-time analysis of nanopore reads with the cloud-based platform (EPI2ME) and its integrated software for sequence similarity-based taxonomic classification (WIMP).

| Sample_Replicate | Yield Data (Mbases) | Average Quality Score | Average Sequence Length | Total Reads | Reads Classified | Superkingdom |
| --- | --- | --- | --- | --- | --- | --- |
| Saliva1_R1 | 33.1 | 11.56 | 734 | 45,079 | 43,820 | Eukaryota: 89%<br>Bacteria: 11%<br>Viruses: < 1% |
| Saliva1_R2 | 25.0 | 11.12 | 847 | 29,462 | 29,059 | Eukaryota: 95%<br>Bacteria: 5%<br>Viruses: < 1% |
| Saliva2_R1 | 223.2 | 12.40 | 729 | 306,111 | 293,219 | Eukaryota: 81%<br>Bacteria: 19%<br>Viruses: < 1%<br>Archaea: < 1% |
| Saliva2_R2 | 66.1 | 10.92 | 672 | 98,330 | 94,121 | Eukaryota: 86%<br>Bacteria: 14%<br>Viruses: < 1% |
| Saliva3_R1 | 22.8 | 11.29 | 497 | 45,827 | 33,609 | Bacteria: 89%<br>Eukaryota: < 11%<br>Viruses: < 1%<br>Archaea: < 1% |
| Saliva3_R2 | 10.7 | 11.47 | 445 | 24,099 | 17,798 | Bacteria: 90%<br>Eukaryota: < 10%<br>Viruses: < 1% |
| Stool1_R1 | 37.3 | 11.70 | 428 | 87,146 | 50,254 | Bacteria: 100%<br>Eukaryota: < 1%<br>Viruses: < 1%<br>Archaea: < 1% |
| Stool1_R2 | 58.3 | 11.63 | 451 | 129,091 | 74,810 | Bacteria: 100%<br>Eukaryota: < 1%<br>Viruses: < 1%<br>Archaea: < 1% |
| Stool2_R1 | 15.9 | 11.21 | 431 | 36,832 | 22,744 | Bacteria: 97%<br>Viruses: 2%<br>Archaea: < 1%<br>Eukaryota: < 1% |
| Stool2_R2 | 21.1 | 11.04 | 566 | 37,172 | 22,425 | Bacteria: 98%<br>Viruses: < 1%<br>Archaea: < 1%<br>Eukaryota: < 1% |

### Slow sequencing shows complex host-microbe interaction types

We investigated the microbial species from these microbiomes further by assigning each microbial species or strain as a harmful, beneficial, or commensal organism in the oral or gut microbiome (Tables S4-5). This assessment of the host-microbe interaction was initially conducted by matching the name of each organism to the list generated by the previous studies to have a positive, negative, or neutral impact on the human host [32,39–45]. The ten most abundant species in each microbiome sample are shown with the host-microbe interaction type as beneficial, harmful, or commensal in Table S4. This curated list shows that the ten most abundant species are consistently present in most replicates of the Saliva samples. For example, the most abundant species of beneficial bacteria are found to be *Haemophilus parainfluenzae* in all the saliva replicates. In contrast, the most abundant species of harmful bacteria are found to be *Neisseria subflava* in Saliva1_R1, whereas *Prevotella melaninogenica* in Saliva1_R2 and Saliva2. In Saliva3, the most abundant harmful bacteria is found to be *Rothia mucilaginosa* in both replicates. Despite the difference in order, the ten most abundant species mostly match between two replicates of the microbiome sample. However, there was much more variation in the ten most abundant species in the Stool samples. For example, the most abundant species of beneficial bacteria is *Lactobacillus ruminis* in both replicates of Stool1, whereas *Akkermansia muciniphila* in Stool2_1. *Bifidobacterium adolescentis* is found in all stool samples as one of the most abundant beneficial bacteria. In both replicates of Stool1, the most abundant harmful bacteria is found to be *Acidaminococcus intestini*, which has been isolated from different clinical samples [46]. In both replicates of Stool2, the most abundant harmful bacteria is found to be *Desulfovibrio piger*, which are sulfate-reducing bacteria that may contribute to gastrointestinal diseases such as inflammatory bowel diseases (IBDs) due to the production of hydrogen sulfide that is toxic to the gut epithelium [47].

Due to the extensive list of microbial species in the nanopore dataset, there were many microbes that were missing from the initial list of host-microbe interaction types, particularly in the gut microbiome which contained thousands of species. Thus, we further searched the most recent scientific literature to assess the impact of each microbial organism in these microbiomes (Table S5). In cases when there is contradicting evidence, we flagged the organisms as inconclusive. Furthermore, if the assessment level was higher than that of the genus, it was immediately assessed as inconclusive (as there is too much diversity) unless there was overwhelming evidence otherwise (Table 2).

**Table 2.**
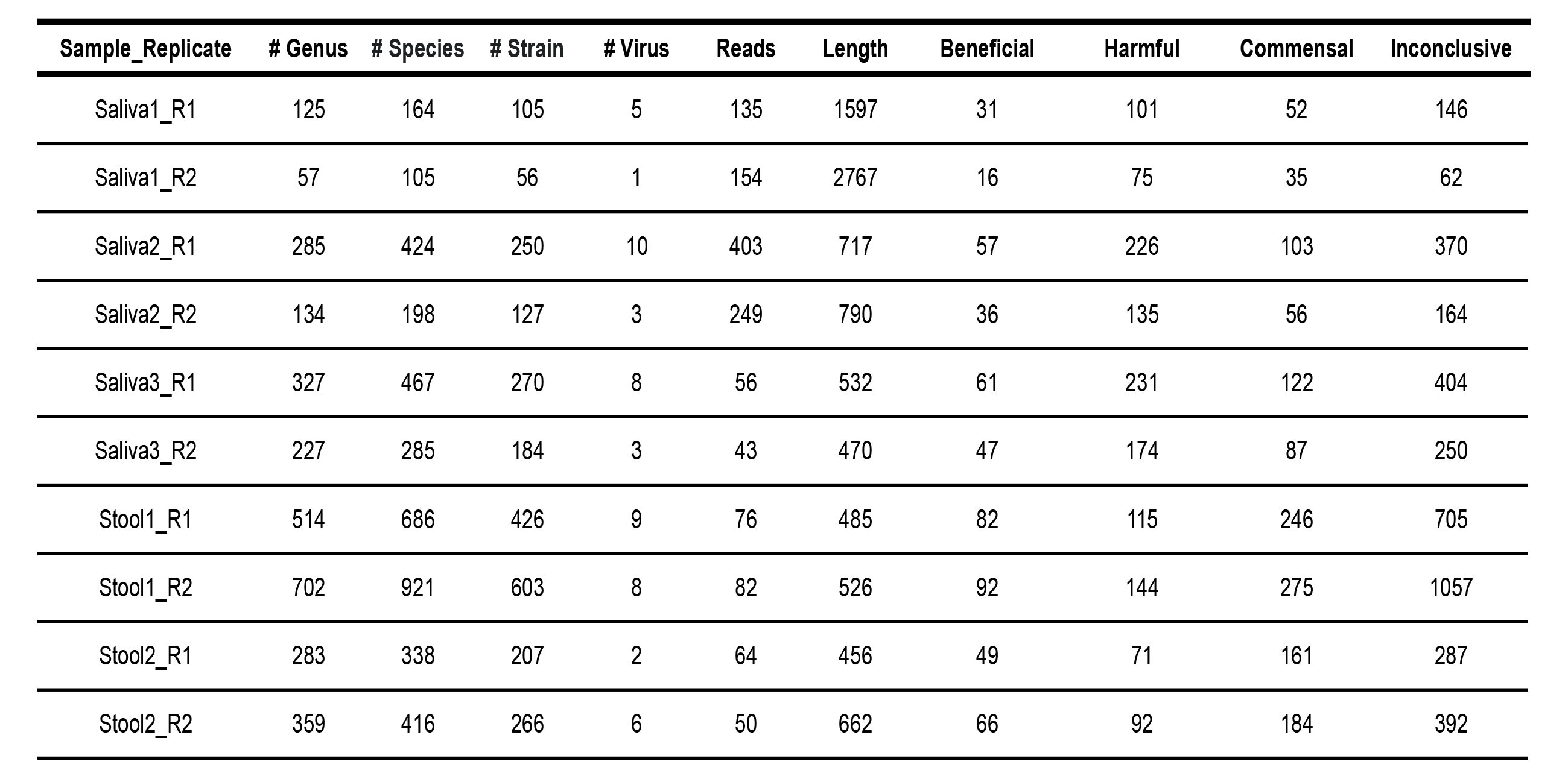
Assessment of host-microbe interaction types for each microbe species per microbiome replicate.

The comprehensive assessment of the host-microbe interaction types in the microbiome community is summarized in the bar chart of relative diversity (Figure 2). The bar chart shows that the oral microbiome tends to contain more diverse organisms that are known to be harmful than the gut microbiome. Moreover, a significant number of microbes exhibit inconclusive roles within the gut and oral microbiomes, underscoring the imperative to explore the impact of these microbes on microbiome communities in order to comprehensively map the landscape of the human microbiome. A bacterial species that have been isolated from a human gut may be beneficial or pathogenic depending on the individual or the health condition of the individual, leading to conflicting or inconclusive information about the host-microbe interaction type. Furthermore, one species may have many strains with completely different characteristics. In our dataset, there are some bacterial species such as *Escherichia coli* with dozens of strains, with a huge diversity in their genomic and functional characteristics. Therefore, even if one sequenced species was considered as one interaction type, there is no guarantee that the actual strain that was sequenced possesses the same interaction type.

**Fig. 2:**
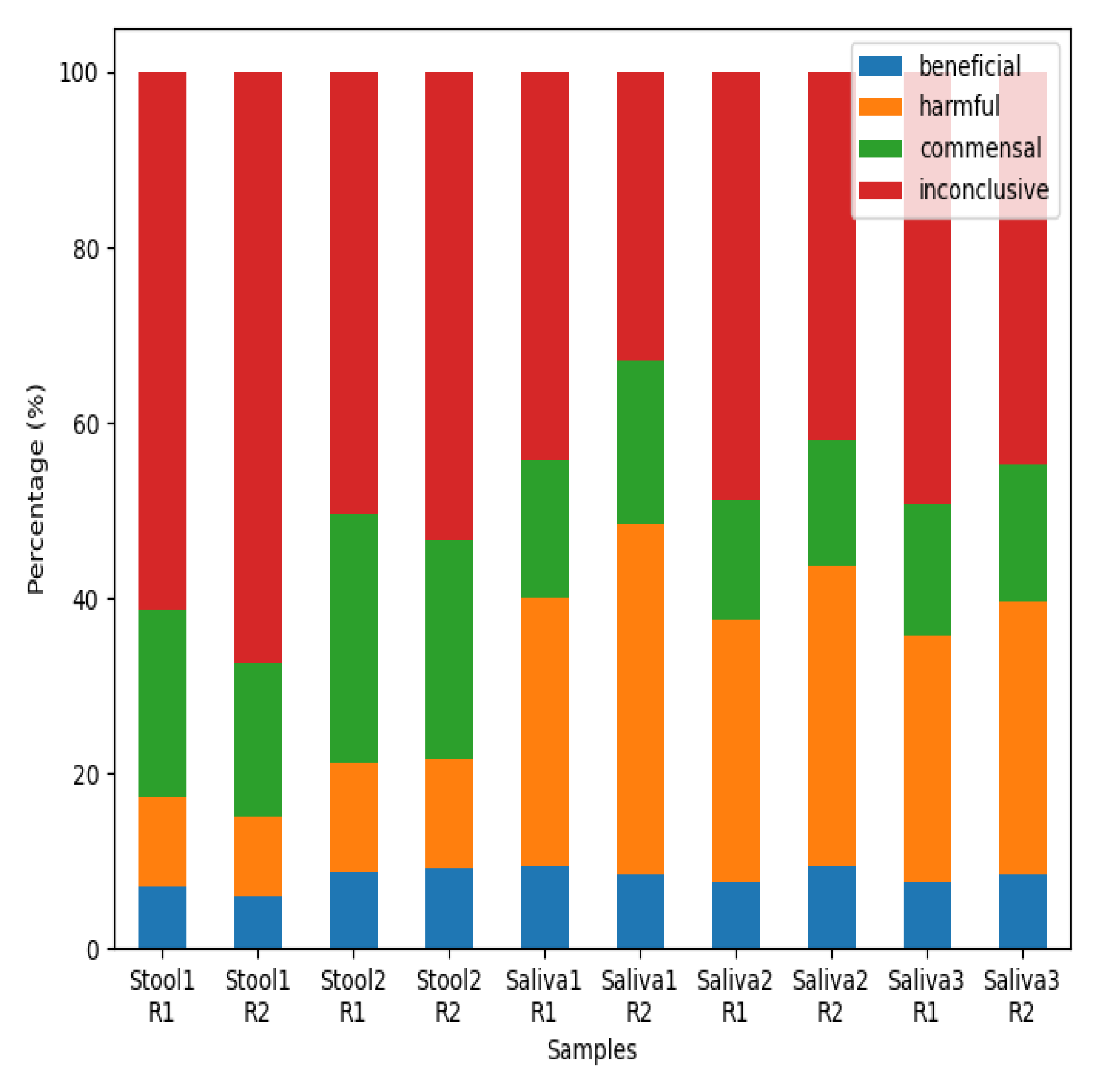
Stacked bar charts of host-microbe interaction types in each microbiome replicate. The stacked bars visually depict the species diversity within each replicate, with classification into beneficial, harmful, or commensal categories in the host microbiome based on existing literature. Any microbial species that have missing or conflicting information is categorized as inconclusive.

We found some microbes whose presence in the gut and oral microbiomes was particularly intriguing (Tables S4 and S5). For example, *Oceanobacillus iheyensis* normally found in the deep sea [48] was found in the oral microbiome of Saliva2_R1. Cellulolytic bacteria (in Caldicellulosiruptor) were sequenced in the microbiomes of Stool1 and Stool2_R2. No evidence was found for its host-microbe interaction type, but cellulolytic bacteria are important for mammals including humans, as they allow the digestion of plant materials and gain nutrients from plants. A previous study even shows the potential for these microbes to have antibacterial properties against pathogenic bacteria [49], which makes it difficult to assess their host-microbe interaction type as commensal or beneficial (list of cellulolytic bacteria includes *Caldicellulosiruptor bescii* DSM 6725, *Caldicellulosiruptor changbaiensis*, *Caldicellulosiruptor obsidiansis* OB47, *Caldicellulosiruptor saccharolyticus* DSM 8903). There are some plant bacteria, both beneficial and pathogenic to plants, whose role in human health has not been investigated. A plant pathogen that is also pathogenic to humans was found in Stool2_R1, such as *Pantoea ananatis*, whose presence is uncommon in the human microbiome [50]. There are some zoonotic bacteria found in the sample, including *Pasteurella multocida* [51]. Some bacteria have a natural affinity towards antimicrobial resistance, including *Clostridium boltae*, which is a commensal in the human gastrointestinal tract but also acts as a reservoir of antimicrobial resistance [52].

During the assessment, we noticed that defining the host-microbe interaction types as harmful, beneficial, or commensal is only a vague indicator of the microbial characteristics, and should not be considered as an absolute measure. For example, *Corynebacterium matruchotii* has been known to cause calcium formation if present in the oral microbiome [53]. The formation could help prevent caries formation but also lead to periodontal diseases, highlighting the dual nature of the host-microbe interaction as both beneficial and harmful. A lot of the microbes that are commensal can also be harmful to immunocompromised patients [54], and the microbial pathogenicity or virulence can undergo changes due to the changes in the microbial DNA, the antimicrobial resistance, the surrounding environment, or the susceptibility of humans to particular diseases [55–57]. For example, *Acinetobacter baumannii* was pathogenic since the 1990s but its pathogenicity level has now increased to a critical level [55]. Furthermore, the composition of the microbiota is as important as the type, as the interplay between different microbes also changes the extent of beneficial or harmful effects [58].

### Oral and gut microbiomes have numerous AMR genes

We found numerous and diverse antibiotic resistance genes in all the microbiome samples, summarized in Table S6 and shown as a heatmap in Figure S4. There are several genes that are attributed to the antimicrobial resistance to a wide range of antibiotics, including beta-lactam, aminoglycoside, tetracycline, macrolide, and fluoroquinolone (Table S7). Bacteria can develop resistance against these antibiotics through multiple mechanisms. These antimicrobial resistance genes can be categorized into four Comprehensive Antibiotic Resistance Database (CARD) models depending on the type of resistance mechanisms: protein variant model, protein homolog model, protein wild type model, and rRNA mutation model (Figure S5).

One of the antibiotics named aminoglycoside is widely used to fight against bacteria, especially aerobic gram-negative bacteria. Aminoglycoside inhibits peptide elongation at 30S ribosomal subunit, resulting in inaccurate mRNA translation which can halt protein synthesis or alter amino acid compositions at certain points [59]. However, when some mutations occur in the 30S ribosomal subunit, aminoglycosides no longer interact with the target [60]. In all the microbiome samples, the AMR genes conferring resistance to aminoglycoside were the most prevalent (Figure 3). For instance, at least 60% of the AMR genes are related to the resistance against aminoglycoside in Saliva3_R2 (Table S6). Particularly, we found *Mycobacterium tuberculosis* in one of the saliva samples (Saliva3_R2), which is known to cause tuberculosis and it had 16s rRNA variant genes that confer multidrug resistance to streptomycin and amikacin, which belong to the family of aminoglycoside, posing a potential threat as these antibiotics are commonly used to treat tuberculosis [61]. In one of the stool samples (Stool1_R1), *Campylobacter jejuni* known to cause gastroenteritis was found. It had ant(6)-Ib genes, which encode a family of aminoglycoside nucleotidyltransferase named ANT(6)-Ib. The expression of ant(6)-Ib can exacerbate the antimicrobial resistance in *Campylobacter jejuni*, as aminoglycosides and macrolides are the effective way to treat this disease [62].

**Fig. 3:**
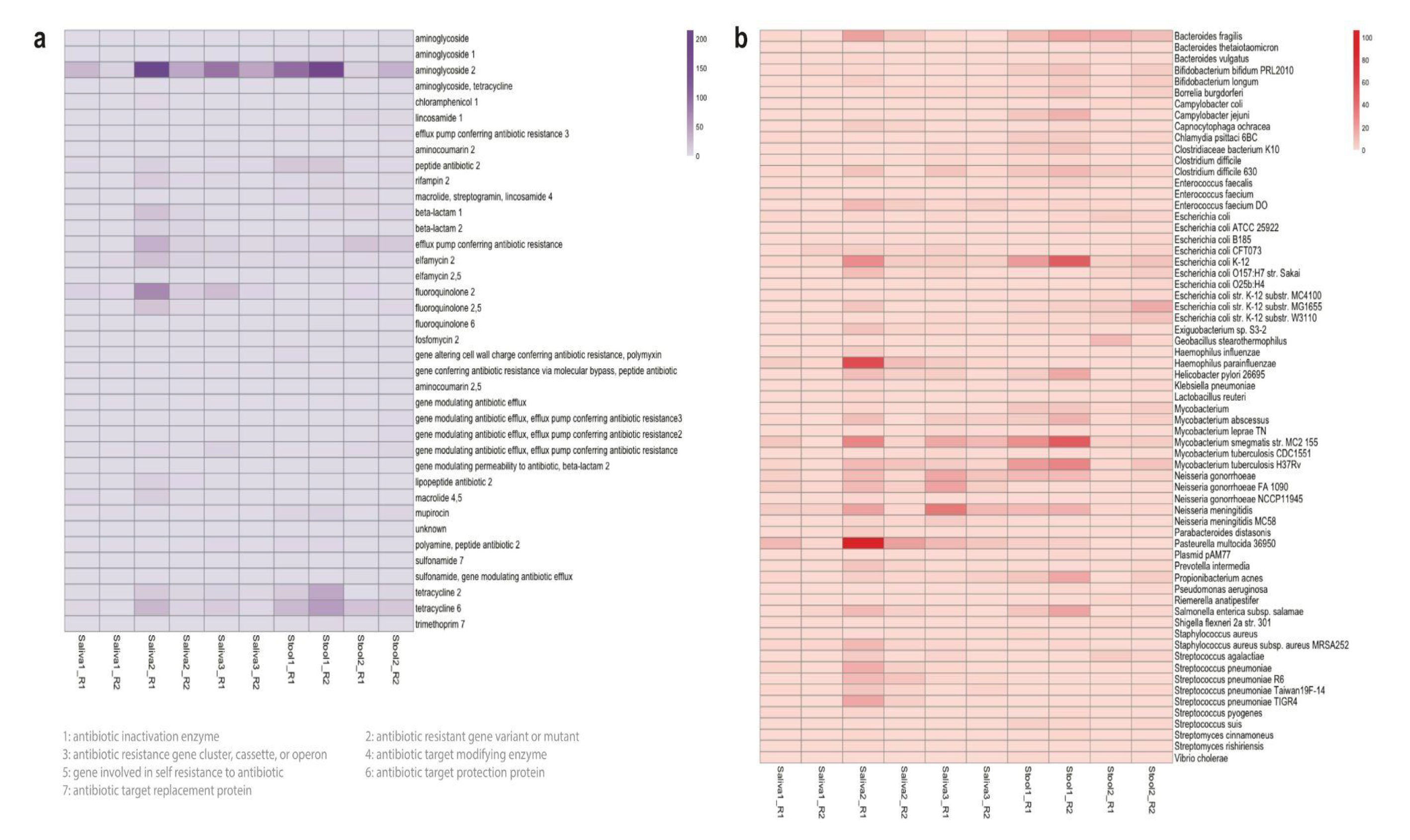
The Comprehensive Antibiotic Resistance Database (CARD) resistance ontology in each microbiome replicate based on (a) antibiotic category and (b) taxon conferring resistance to antibiotics. Antibiotics are classified based on their mechanism of action, spectrum of activity, or chemical structure. The antibiotic category shows all resistance pathways linking the gene to antibiotic molecules.

As shown in our microbiome samples, many conventional antibiotics as well as some newer antibiotics are no longer effective in certain types of bacteria due to the spread of antimicrobial resistance. Recently, the World Health Organization (WHO) has designated antimicrobial resistance as one of the top threats against public health and published a list of pathogens that are in urgent need of novel antibiotics [2]. The WHO list is divided into three levels of priority (critical, high, and medium) according to the severity of antimicrobial resistance and the urgency for novel antibiotics. We compared the WHO list with the microbial species present in each microbiome sample, and we found three bacterial species (*Neisseria gonorrhoeae*, *Shigella flexneri*, *Streptococcus pneumoniae*) that matched (Table 3). In all the saliva samples, we found *Neisseria gonorrhoeae* included in the high-priority category (Table 3), which are found to be resistant to cephalosporin or fluoroquinolone (Table S5). Fluoroquinolones are one of the most important antibiotics listed by the WHO, as they inhibit DNA replication by preventing the ligase activity of the bacterial DNA gyrase and topoisomerase IV [63]. In gram-negative bacteria, plasmid-mediated resistance genes produce proteins that can bind to the bacterial DNA gyrase, protecting it from the action of quinolones [64].

**Table 3.** The AMR-conferring taxa and their characteristics in the oral and gut microbiome of the human samples. Multidrug therapy implies that this pathogen requires multiple antibiotics to treat the related disease.

| Gram-Positive |  |  |  |  |  |  |
| --- | --- | --- | --- | --- | --- | --- |
| Phylum | Taxon | Colony | Spore | Respiration | Disease | Antimicrobial therapy |
| Actinomycetota | *Bifidobacterium bifidum* | Rod | — | Anaerobic | — | — |
| Actinomycetota | *Bifidobacterium longum* | Rod | — | Anaerobic | — | — |
| Actinomycetota | *Cutibacterium acnes* | Rod | — | Anaerobic | Skin infections | Benzoyl peroxide |
| Actinomycetota | *Mycobacterium leprae* | Rod | — | Aerobic | Hansen's disease | Multidrug therapy |
| Actinomycetota | *Mycobacterium smegmatis* | Rod | — | Aerobic | — | — |
| Actinomycetota | *Mycobacterium tuberculosis* | Rod | — | Aerobic | Tuberculosis | Multidrug therapy |
| Actinomycetota | *Mycobacteroides abscessus* | Rod | — | Aerobic | Lung disease | Macrolide |
| Actinomycetota | *Mycobacteroides chelonae* | Rod | — | Aerobic | Skin infections | Macrolide |
| Actinomycetota | *Streptomyces cinnamoneus* | Filamentous | ○ | Aerobic | — | — |
| Actinomycetota | *Streptomyces rishiriensis* | Filamentous | ○ | Aerobic | — | — |
| Bacillota | *Clostridioides difficile* | Rod | ○ | Anaerobic | Colon infections | Glycopeptide |
| Bacillota | *Enterococcus faecium* | Cocci | — | Facultative anaerobic | Urinary tract infections | Glycopeptide |
| Bacillota | *Enterococcus faecium* | Rod | — | Facultative anaerobic | — | — |
| Bacillota | *Lactobacillus reuteri* | Rod | — | Anaerobic | — | — |
| Bacillota | *Staphylococcus aureus* | Cocci | — | Facultative anaerobic | Skin infections | Oxazolidinone |
| Bacillota | *Streptococcus agalactiae* | Cocci | — | Facultative anaerobic | Group B Streptococcal (GBS) infections | Ampicillin |
| Bacillota | *Streptococcus pneumoniae* | Diplococci | — | Facultative anaerobic | Pneumonia | Multidrug therapy |
| Bacillota | *Streptococcus pyogenes* | Cocci | — | Facultative anaerobic | Group A Streptococcal (GAS) Infections | Amoxicillin |
| Bacillota | *Streptococcus suis* | Cocci | — | Facultative anaerobic | Zoonotic disease | Aminopenicillin |

Gram-Negative
| Phylum | Taxon | Colony | Spore | Respiration | Disease | Antimicrobial therapy |
| --- | --- | --- | --- | --- | --- | --- |
| Bacteroidota | *Bacteroides fragilis* | Rod | — | Anaerobic | Inflammatory bowel disease | Nitroimidazole |
| Bacteroidota | *Bacteroides vulgatus* | Rod | — | Anaerobic | Inflammatory bowel disease | Nitroimidazole |
| Bacteroidota | *Capnocytophaga ochracea* | Rod | — | Facultative anaerobic | Capnocytophaga infection | Multidrug therapy |
| Bacteroidota | *Parabacteroides distasonis* | Rod | — | Anaerobic | — | — |
| Bacteroidota | *Prevotella intermedia* | Rod | — | Anaerobic | Periodontal infections | Nitroimidazole |
| Campylobacterota | *Campylobacter jejuni* | Rod | — | Microaerophilic | Gastroenteritis | Macrolide |
| Campylobacterota | *Helicobacter pylori* | Helical | — | Microaerophilic | Stomach ulcers | Multidrug therapy |
| Chlamydiota | *Chlamydia psittaci* | Cocci | — | Anaerobic | Psittacosis | Macrolide |
| Pseudomonadota | *Escherichia coli* | Rod | — | Facultative anaerobic | Escherichia coli infection | Tetracycline |
| Pseudomonadota | *Haemophilus parainfluenzae* | Cocci | — | Facultative anaerobic | Pneumonia | Cephalosporin |
| Pseudomonadota | *Klebsiella pneumoniae* | Rod | — | Facultative anaerobic | Klebsiella pneumoniae infection | Carbapenem |
| Pseudomonadota | *Neisseria gonorrhoeae* | Diplococci | — | Anaerobic | Gonorrhea | Cephalosporin |
| Pseudomonadota | *Neisseria meningitidis* | Diplococci | — | Anaerobic | Meningitis | Cephalosporin |
| Pseudomonadota | *Pasteurella multocida* | Cocci | — | Facultative anaerobic | Subcutaneous infection | Aminopenicillin |
| Pseudomonadota | *Salmonella enterica* | Rod | — | Facultative anaerobic | Salmonellosis | Fluoroquinolone |
| Pseudomonadota | *Shigella flexneri* | Rod | — | Facultative anaerobic | Shigellosis | Fluoroquinolone |
| Pseudomonadota | *Vibrio cholerae* | Rod | — | Facultative anaerobic | Cholera infection | Tetracycline |
| Spirochaetota | *Borrelia burgdorferi* | Helical | — | Anaerobic | Lyme disease | Tetracycline |

In some microbiome samples, we found some bacteria of a medium priority category from the WHO list, including *Shigella* which are also resistant to fluoroquinolone (Table S5). One of the stool samples (Stool2_R1) contains the same genus of bacteria named *Shigella flexneri* (Table 3). The point mutations in the DNA gyrase (gyrA) give rise to fluoroquinolone resistance, and we found the gyrA genes that confer resistance to fluoroquinolone in these bacteria (Table S6). It is intriguing to observe that the saliva and stool microbiomes all had these genes because they are known to cause cross-resistance to fluoroquinolones. For instance, recent research shows the *Mycobacterium tuberculosis* strain with the gene variant gyrA exhibits cross-resistance to six different fluoroquinolones, whereas the strain which does not have mutations in gyrA shows resistance specifically to the particular fluoroquinolones [65]. Another bacteria that matched the medium priority category is *Streptococcus pneumoniae* (Table 3), which is no longer susceptible to penicillin (Table S5). Bacteria can acquire resistance by synthesizing an enzyme such as beta-lactamase that attacks the beta-lactam ring of penicillin molecules. There are also other ways to become penicillin-resistant through mechanisms that decrease the binding affinity of the antibiotics. In all the saliva microbiome samples, *Streptococcus pneumoniae* has mutated variants of PBP1a, PBP2b, and PBP2x (Table S4). These penicillin-binding proteins (PBPs) are targeted by beta-lactam antibiotics [66], thus these mutations in the PBPs can lead to resistance against penicillin.

In the AMR analysis, we noticed that a wide variety of nonpathogenic bacteria have numerous and diverse AMR-related genes (Figure 3b). For example, *Mycobacterium smegmatis* are nonpathogenic bacteria but they are one of the most abundant bacteria that confer resistance to antibiotics in both the oral and gut microbiomes. *Haemophilus parainfluenzae* and *Bacteroides fragilis* are other examples of nonpathogenic bacteria that are present across all the microbiomes. These nonpathogenic bacteria are potential reservoirs for AMR-related genes through horizontal gene transfer, which is the primary mechanism for the spread of antibiotic resistance in bacteria [67]. Nonpathogenic bacteria which are in the same genus as pathogenic bacteria are of particular concern as their horizontal gene transfer is facilitated. For example, both *Mycobacterium tuberculosis* and *Mycobacterium smegmatis* with AMR-related genes are present in both the oral and gut microbiomes with high abundance [68].

### Deep learning-based classification of unidentified microbes predicts mobilome

The fast sequencing mode of the nanopore data involves the taxonomic classification of metagenomic sequences in real-time. This fast mode is enabled by a cloud-based platform integrated into the sequencing software, and it utilizes the benefits of long reads to enable rapid species identification and quantification from metagenomic samples based on the sequence similarity algorithm [29]. However, this sequence similarity-based approach does not fully exploit the potential of nanopore sequencing to produce long-read DNAs that can be regarded as a long stretch of DNA from a microbe, or even an individual.

During the fast sequencing analysis, we noticed that there were many ‘unclassified’ reads in the classification results based on the sequence similarity algorithm. On average, the oral microbiome had around 10,000 unclassified reads and the gut microbiome had around 40,000 unclassified reads. We assumed that these unclassified reads are unidentifiable as unexplored organisms in the human microbiome, and we had a hypothesis that many of these unidentified reads are from mobile genetic elements such as bacteriophages and plasmids.

To test this hypothesis that these unidentified reads derive from mobile genetic elements, we searched for a different type of taxonomic algorithm that can classify a sequence without the presence of similar sequences in the search database. We found that deep learning-based algorithms can place de novo sequences in taxonomic categories with high accuracy when trained with a huge diversity and quantity of genetic sequences, exploiting the fact that different species have their specific patterns and characteristics engraved in their genetic information [69]. For example, eukaryotic genomes tend to have more noncoding regions compared to prokaryotic genomes, whereas bacteriophages are recently found to adapt alternative genetic coding to increase fitness and evolvability [70,71].

In this slow sequencing mode, we analyzed each unidentified read using a deep learning-based approach to assign taxonomic classification at the superkingdom level (Figure 4). The heatmaps show the predicted phylum of all the samples for each superkingdom, revealing the stool samples have more diversity in the four superkingdoms than the saliva samples as expected (Figure S5). The heatmap of the virus superkingdom is of particular interest, which is labeled with the predicted host phylum of each read. According to the deep learning-based approach, the oral and gut microbiomes are expected to have diverse viruses against archaea, bacteria, and eukaryotes, including against Actinobacteria, Crenarchaeota, and Arthropoda. Another interesting observation is that many DNA reads are still unknown even after the deep learning-based classification that does not utilize any database for inference. This reveals that some de novo reads in these microbiomes are completely devoid of any known patterns and characteristics, which is an intriguing observation to be investigated further.

**Fig. 4:**
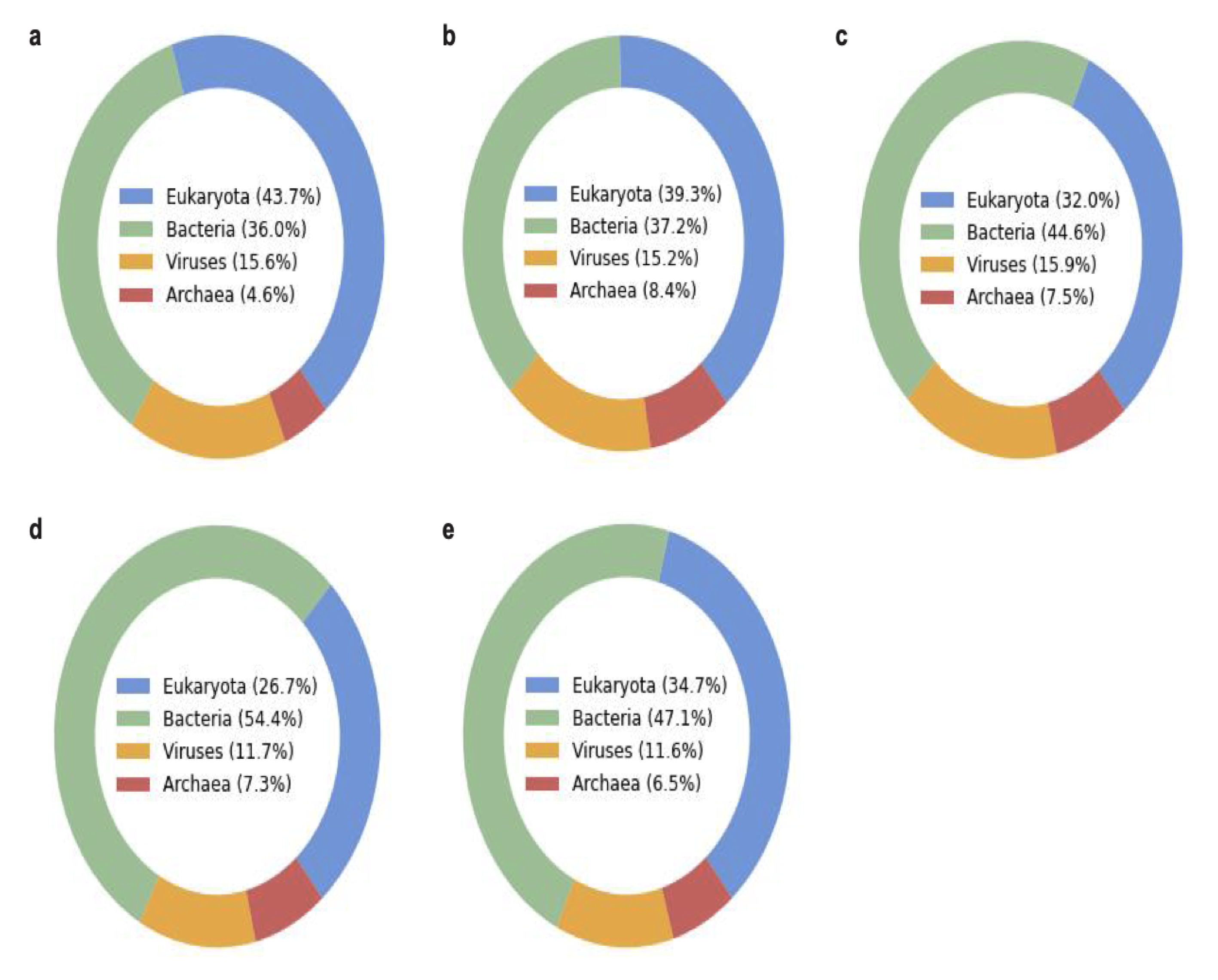
Superkingdom of unidentified reads predicted by the deep learning-based algorithm (BERTax) in each microbiome sample (a) Saliva1, (b) Saliva2, (c) Saliva3, (d) Stool1, and (e) Stool2. The two replicates per sample were combined for this exploratory analysis.

Followingly, the predicted classification of unidentified reads from each microbiome sample is separated into four superkingdoms of archaea, bacteria, eukaryotes, and viruses, and summarized into the bar charts at the genus level (Figure S6-S11). The deep learning-based classification of this dataset at the genus level shows an intriguing range of diversity in the classification. Particularly, the diversity at the genus level of the eukaryotic organisms was rich in all the microbiome samples, but this may be due to the training dataset of the deep learning-based approach having a bias towards eukaryotic genomes [69]. The statical summary of this analysis shows that many of the unidentified reads are classified into the virus category according to the deep learning-based algorithm (Table S6). This number is overrepresented as compared to the previous taxonomic classification of viruses based on sequence similarity (Table S3). The bar chart of the predicted virus at the genus level (Figure S11) is particularly interesting as they reveal the unexplored territory of mobilome in the human microbiome that is yet to be discovered for novel therapeutic tools and bacteriophage therapy [72–75]. We noticed some inconsistencies in the prediction at the different levels of superkingdom, phylum, and genus, thus this deep learning-based approach should be regarded more as an exploratory tool rather than a diagnostic tool.

## Discussion

The development and implementation of rapid and accurate diagnostic tools for bacterial infections are essential in combating the current crisis of antimicrobial resistance (AMR) effectively. This requires a shift away from traditional culture-based techniques towards molecular-based diagnostic methods, which can provide faster and more accurate results, leading to better patient outcomes. Here, we focused on the ability of nanopore sequencing to generate long-read native DNAs from metagenomic samples of various human microbiomes. Nanopore sequencing enables direct analysis of DNA/RNA sequences by sensing changes in an electric current as they pass through a protein nanopore [76]. This new sequencing technology is revolutionizing genomics, as it can produce long-read DNA/RNA sequences allowing genomic analysis of microbes at individual levels. We explored the potential of nanopore sequencing as a futuristic diagnostic tool, which could provide ultrarapid profiling of the human microbiome through real-time analysis such as species identification and antimicrobial resistance [2].

We further explored the potential of nanopore sequencing to be utilized in two different modes as a diagnostic tool: fast sequencing and slow sequencing. The fast mode enables real-time analysis of pathogen identification, metagenomic analysis of microbial communities, and antimicrobial resistance analysis. This mode is rapid and direct, requiring minimal inputs of human expertise and curation. In this fast analysis, we classified thousands of microbial species in the saliva and stool samples, with the most cost-effective but a lower-yield and single-use version of nanopore flow cells [77]. Furthermore, we rapidly identified the ten most abundant species that are known to be beneficial, harmful, or commensal in the oral and gut microbiome using the previously curated list. The slow mode enables in-depth analysis of host-microbe interactions and deep learning-based classification of unidentified reads. This mode is deliberate and exploratory, requiring the most advanced bioinformatic skills and expertise in microbiome research. A thorough exploration of host-microbe interaction types underscores the existing knowledge gaps regarding the impact of numerous microbes identified within the oral and gut microbiomes. Additionally, we evaluated a largely unexplored dataset of unclassified DNA reads from the sequence similarity-based analysis by utilizing a deep learning-based algorithm that does not require a match in the database to predict the superkingdom, phylum, and genus of these reads. The analysis further uncovers the potential existence of diverse organisms belonging to bacteria, archaea, and eukaryotes, with a significantly higher proportion of reads predicted to originate from virus genomes.

In this study, we aim to provide an exploratory application of nanopore sequencing as a future diagnostic tool for bacterial infection, which has resurfaced in the scientific community as an urgent global health issue due to the uncontrolled spread of antimicrobial resistance [78]. Nevertheless, it is important to acknowledge several caveats that were encountered during this exploratory application. Firstly, there are still debates about the accuracy of nanopore sequencers at simplex sequencing, which depends on the nanopore version, chemistry, and basecalling algorithms. According to the manufacturer, we used the flow cell version and chemistry (R9.4.1 and SQK-LSK110, respectively) with the expected raw-read accuracy of 98.3% modal. Regarding the accuracy of read classification, a recent study investigated that the taxonomic classification of long-read DNAs is satisfactory through controlled experiments using mock microbial communities [79]. This study further demonstrated that the expected microbial species corresponded at anticipated abundances, with the limit of detection observed at 4 reads and 5,000 bp in length. However, we still had difficulties in determining the confidence level of very rare species, despite setting a high Q-score threshold while using the high-accuracy basecalling. Since the current state-of-the-art sequencing technologies cannot provide ground truth about the presence of these rare species in metagenomic samples, we utilized two replicates per sample to build confidence in the results of species identification.

Secondly, we used a specialized microbiome kit for extracting native DNA from different human microbiomes. While the kit was successful in extracting sufficient amounts of DNA from saliva and stool samples, it was not effective for extracting DNA from urine, nasal, and vaginal samples (Table S1). Alternative extraction methods that can extract small quantities of microbial DNA more efficiently may be necessary for these types of samples. Finally, the extracted DNA from some saliva samples using this kit still had a substantial fraction of human DNA despite having a host DNA depletion step. We suggest using other methods of human DNA depletion to enrich the microbiome DNA against the human DNA [80]. Adaptive sampling has emerged as a cutting-edge approach for selectively reducing host DNA content in human samples [81]. Adaptive sampling is a technique in nanopore sequencing that allows for selective sequencing of specific genomic regions of interest, optimizing the sequencing process by focusing on relevant regions and reducing sequencing time and cost [82]. It involves real-time analysis of the sequencing data and adjustment of the sequencing parameters to increase the coverage of targeted regions.

In conclusion, rapid and accurate pathogen identification and microbial profiling are essential in combating infectious diseases effectively, and the development of new technologies, such as nanopore sequencing, offers great promise as innovative diagnostic tools. The main advantages of nanopore sequencing as a diagnostic tool include a cost-effective sequencer ($1,000) and flow cell ($100 per sample) and flexible adaptation of downstream analysis as a fast mode (< 12 hours to pathogen identification) and a slow mode (several weeks) depending on the type of information needed. Nevertheless, addressing the existing challenges and ensuring the extensive utilization of these technologies in clinical settings necessitates further efforts and advancements.

## Methods and Materials

### Preparation of non-invasion human microbiome sample

Human microbiomes were collected from the healthy volunteers after signing a written informed consent between March 2022 and July 2022. For the saliva samples (Saliva1_R1, Saliva1_R2, Saliva2_R1, Saliva2_R2, Saliva3_R1, Saliva3_R2), saliva collected with sterile medical swabs were transferred to 1000 µl of PBS solution (P5119; Sigma-Aldrich, Darmstadt, Germany). For the stool samples (Stool1_R1, Stool1_R2, Stool2_R1, Stool2_R2), stool collected with sterile medical swabs was transferred to 1000 µl of PBS solution. For the urine, nasal, and vaginal samples (Urine1_R1, Urine1_R2, Nasal1_R1, Nasal1_R2, Vaginal1_R1, Vaginal1_R2), each sample was collected with sterile medical swabs and transferred to 1000 µl of PBS solution.

### Microbiome DNA extraction and quality control

Microbiome DNAs were extracted from the human samples on the same day of collection using the microbiome-specific kit (QIAamp DNA Microbiome Kit; Qiagen, Hilden, Germany). The DNA extraction was performed according to the manufacturer’s protocol. The only modification to the protocol was to conduct all the centrifuge steps at the speed of 12,300 x g instead of 20,000 x g. The extracted DNA samples were quantified for the quantity (ng/µl) and quality (A260/A280 and A260/A230) using a spectrophotometer (NanoDrop 2000; Thermo Scientific, Waltham, USA). The A260/A280 acceptable ratio was kept at 1.8–2.0, and the A260/A230 acceptable ratio was kept at 2.0–2.2 for the quality control for nanopore sequencing (Table S1). The quality-controlled DNA samples were kept at 4 °C until further treatment or analysis was performed.

### Preparation of sequencing library using native DNA ligation

The sequencing library was prepared from at least 500 ng of high molecular weight genomic DNA extracted from the human microbiome samples using the native DNA ligation kit (SQK-LSK110; Oxford Nanopore Technologies, Oxford, UK) according to the manufacturer’s protocol. For Flongle flow cells, an expansion kit (EXP-FSE001; Oxford Nanopore Technologies, Oxford, UK) was additionally needed to prepare the sequencing mix. The NEBNext FFPE Repair Mix (M6630) and NEBNext Ultra II End repair/dA-tailing Module (E6056) reagents were prepared in accordance with the manufacturer’s instructions. The sample purification was performed using magnetic beads (Agencourt AMPure XP; Beckman Coulter, Orange County, USA) and a magnetic separator (DynaMag^TM^-2 Magnet; Thermo Fischer Scientific, Waltham, USA).

### Nanopore sequencing using MinION and Flongle adapter and flow cell

A Flongle flow cell (FLO-FLG001; Oxford Nanopore Technologies, Oxford, UK) was used for each sample, which was inserted into the Nanopore MinION sequencer (Mk1B MIN-101B; Oxford Nanopore Technologies, Oxford, UK) using the Flongle adapter (ADP-FLG001; Oxford Nanopore Technologies, Oxford, UK). Flongle flow cells were first checked for the minimum number of pores (at least 50 pores) before being primed with 119 µl of the priming mix prepared in accordance with the manufacturer’s instructions. In the priming step, some liquid was left in the P200 pipette tip to ensure no air bubble was inserted. 29 µl of the sequencing mix was loaded onto the Flongle flow cell immediately afterward, following the manufacturer’s protocol. Finally, Flongle flow cells were sealed using the adhesive on the seal tab and the platform lid, and nanopore sequencing was performed for at least 12 hours to obtain a maximum read output.

### Real-time high-accuracy basecalling and cloud-based EPI2ME analysis

The MinKNOW software (v.4.5.4; Oxford Nanopore Technologies, Oxford, UK) was used for raw data acquisition. The raw signal data in FAST5 files were basecalled real-time into the DNA reads in FASTQ files using the high-accuracy mode of the Guppy basecaller (v.5.1.13), integrated within the MinKNOW software.

For the rapid downstream analysis, a cloud-based analysis platform providing rapid analysis workflows called EPI2ME was used. Using the EPI2ME platform (v.3.5.7; Oxford Nanopore Technologies, Oxford, UK), species identification with the WIMP workflow (v.2021.11.26) such as fungi, bacteria, viruses, or archaea, was conducted in real-time based on the Centrifuge classification engine [29,83]. Next, antimicrobial resistance analysis was conducted in real-time with the ARMA workflow (v.2021.11.26.) to identify the genes responsible for antibiotic resistance in the DNA reads, based on the CARD database.

### In-depth microbiome analysis of classified reads

The WIMP workflow utilizes long reads from nanopore sequencing to rapidly identify and quantify microbial species from metagenomic samples. The WIMP results from each sample were downloaded as CSV files, which were processed into classified and unclassified categories. The classified reads from the WIMP workflow were saved separately, and the identified species were further categorized into four host-microbe interaction types (beneficial, commensal, harmful, and inconclusive). The initial list of host-microbe interaction types for several microbial species was curated by pooling a number of studies on the oral microbiome [32,39–41] and the gut microbiome [43,44]. However, many of the microbial species in the oral and gut microbiomes were missing from this curated list, which further required an extensive literature review on each microbial species to determine the host-microbe interaction type. When assessing these bacteria into different interaction types, the exact region in the human body was considered. For example, a commensal in the human gut may be assessed as a pathogen in the human skin.

### In-depth microbiome analysis of unclassified reads

The unclassified reads from the WIMP workflow based on the sequence similarity search were saved separately and analyzed with other methods. These latest algorithms for species identification include the BERTax taxonomic classification [69]. The BERTax taxonomic classification is a deep learning approach based on natural language processing [84] to classify the superkingdom and phylum of DNA sequences taxonomically. It achieves the assignment of unknown sequences to biological clades with shared ancestry in data-dependent training without the need for a genome similarity search of large genome databases. BERTax was shown to perform comparably to the state-of-the-art methods for sequences with close relatives in the database and superior for new species [69]. The unclassified reads from the human microbiome samples were run with the BERTax algorithm to assign the superkingdom, phylum, and genus given the patterns of DNA sequences.

## Data Availability

All the nanopore sequences are available on our project GitHub page (https://github.com/hshimlab/Nanopore_microbiome).

## Code Availability

All codes related to this project are available under an open-source license at https://github.com/hshimlab/Nanopore_microbiome. For data analysis, Python .3.6.4 (https://www.python.org), NumPy v.1.17.5 (https://github.com/numpy/numpy), SciPy v.1.1.0 (https://www.scipy.org), seaborn v.0.9.0 (https://github.com/mwaskom/seaborn), Matplotlib v.3.3.4 (https://github.com/matplotlib/matplotlib), pandas v.0.22.0 (https://github.com/pandas-dev/pandas) were used. For nanopore data acquisition, we used the MinKNOW v.21.11.8 and MinKNOW core v.4.5.4. For rapid nanopore data analysis, we used the EPI2ME platform v.3.5.7. For taxonomic analysis, we used the BERTax taxonomic classification (https://github.com/f-kretschmer/bertax).

## Acknowledgments (optional)

We thank the members of the Center for Biotech Data Science at GUGC for encouragement, support, and motivation. The research and development activities described in this study were funded by Ghent University Global Campus (GUGC), Incheon, Korea. We would like to acknowledge Daniel Kahneman, the author of ‘Thinking, Fast and Slow’ for inspiring the concept of two systems of sequencing.

## Competing interests

None

## Supplementary Information

Supplementary Information will be provided upon request.

## Notes

### Competing Interest Statement

The authors have declared no competing interest.

## References

1. Maurer FP, Christner M, Hentschke M, Rohde H. Advances in Rapid Identification and Susceptibility Testing of Bacteria in the Clinical Microbiology Laboratory: Implications for Patient Care and Antimicrobial Stewardship Programs. Infect Dis Rep. 2017;9: 6839.

2. Shim H. Three Innovations of Next-Generation Antibiotics: Evolvability, Specificity, and Non-Immunogenicity. Antibiotics (Basel). 2023;12. doi:10.3390/antibiotics12020204

3. Tsalik EL, Petzold E, Kreiswirth BN, Bonomo RA, Banerjee R, Lautenbach E, et al. Advancing Diagnostics to Address Antibacterial Resistance: The Diagnostics and Devices Committee of the Antibacterial Resistance Leadership Group. Clin Infect Dis. 2017;64: S41–S47.

4. Preidis GA, Versalovic J. Targeting the human microbiome with antibiotics, probiotics, and prebiotics: gastroenterology enters the metagenomics era. Gastroenterology. 2009;136: 2015–2031.

5. Frank DN, St Amand AL, Feldman RA, Boedeker EC, Harpaz N, Pace NR. Molecular-phylogenetic characterization of microbial community imbalances in human inflammatory bowel diseases. Proc Natl Acad Sci U S A. 2007;104: 13780–13785.

6. Gonzalez A, Stombaugh J, Lozupone C, Turnbaugh PJ, Gordon JI, Knight R. The mind-body-microbial continuum. Dialogues Clin Neurosci. 2011;13: 55–62.

7. Belkaid Y, Hand TW. Role of the microbiota in immunity and inflammation. Cell. 2014;157: 121–141.

8. Buffie CG, Pamer EG. Microbiota-mediated colonization resistance against intestinal pathogens. Nat Rev Immunol. 2013;13: 790–801.

9. Caballero S, Pamer EG. Microbiota-mediated inflammation and antimicrobial defense in the intestine. Annu Rev Immunol. 2015;33: 227–256.

10. Sommer F, Bäckhed F. The gut microbiota--masters of host development and physiology. Nat Rev Microbiol. 2013;11: 227–238.

11. O’Toole PW, Jeffery IB. Gut microbiota and aging. Science. 2015;350: 1214–1215.

12. Barczak AK, Gomez JE, Kaufmann BB, Hinson ER, Cosimi L, Borowsky ML, et al. RNA signatures allow rapid identification of pathogens and antibiotic susceptibilities. Proc Natl Acad Sci U S A. 2012;109: 6217–6222.

13. Caliendo AM, Gilbert DN, Ginocchio CC, Hanson KE, May L, Quinn TC, et al. Better tests, better care: improved diagnostics for infectious diseases. Clin Infect Dis. 2013;57 Suppl 3. doi:10.1093/cid/cit578

14. Salipante SJ, Sengupta DJ, Rosenthal C, Costa G, Spangler J, Sims EH, et al. Rapid 16S rRNA next-generation sequencing of polymicrobial clinical samples for diagnosis of complex bacterial infections. PLoS One. 2013;8: e65226.

15. Bradley P, Gordon NC, Walker TM, Dunn L, Heys S, Huang B, et al. Rapid antibiotic-resistance predictions from genome sequence data for Staphylococcus aureus and Mycobacterium tuberculosis. Nat Commun. 2015;6: 10063.

16. Ayling M, Clark MD, Leggett RM. New approaches for metagenome assembly with short reads. Brief Bioinform. 2020;21: 584–594.

17. Branton D, Deamer DW, Marziali A, Bayley H, Benner SA, Butler T, et al. The potential and challenges of nanopore sequencing. Nat Biotechnol. 2008;26: 1146–1153.

18. Quick J, Quinlan AR, Loman NJ. A reference bacterial genome dataset generated on the MinION^TM^ portable single-molecule nanopore sequencer. GigaScience. 2014. doi:10.1186/2047-217x-3-22

19. Niedringhaus TP, Milanova D, Kerby MB, Snyder MP, Barron AE. Landscape of next-generation sequencing technologies. Anal Chem. 2011;83: 4327–4341.

20. Latorre-Pérez A, Villalba-Bermell P, Pascual J, Vilanova C. Assembly methods for nanopore-based metagenomic sequencing: a comparative study. Sci Rep. 2020;10: 13588.

21. Charalampous T, Kay GL, Richardson H, Aydin A, Baldan R, Jeanes C, et al. Nanopore metagenomics enables rapid clinical diagnosis of bacterial lower respiratory infection. Nat Biotechnol. 2019;37: 783–792.

22. Petersen LM, Martin IW, Moschetti WE, Kershaw CM, Tsongalis GJ. Third-Generation Sequencing in the Clinical Laboratory: Exploring the Advantages and Challenges of Nanopore Sequencing. J Clin Microbiol. 2019;58. doi:10.1128/JCM.01315-19

23. MacKenzie M, Argyropoulos C. An Introduction to Nanopore Sequencing: Past, Present, and Future Considerations. Micromachines (Basel). 2023;14. doi:10.3390/mi14020459

24. Jain M, Koren S, Miga KH, Quick J, Rand AC, Sasani TA, et al. Nanopore sequencing and assembly of a human genome with ultra-long reads. Nat Biotechnol. 2018;36: 338–345.

25. Vereecke N, Bokma J, Haesebrouck F, Nauwynck H, Boyen F, Pardon B, et al. High quality genome assemblies of Mycoplasma bovis using a taxon-specific Bonito basecaller for MinION and Flongle long-read nanopore sequencing. BMC Bioinformatics. 2020;21. doi:10.1186/s12859-020-03856-0

26. Napit R, Manandhar P, Chaudhary A, Shrestha B, Poudel A, Raut R, et al. Rapid genomic surveillance of SARS-CoV-2 in a dense urban community of Kathmandu Valley using sewage samples. PLoS One. 2023;18: e0283664.

27. Ewing B, Hillier L, Wendl MC, Green P. Base-calling of automated sequencer traces using phred. I. Accuracy assessment. Genome Res. 1998;8: 175–185.

28. Delahaye C, Nicolas J. Sequencing DNA with nanopores: Troubles and biases. PLoS One. 2021;16: e0257521.

29. Kim D, Song L, Breitwieser FP, Salzberg SL. Centrifuge: rapid and sensitive classification of metagenomic sequences. Genome Res. 2016;26: 1721.

30. O’ Donnell MM, Harris HMB, Lynch DB, Ross RP, O’Toole PW. Lactobacillus ruminis strains cluster according to their mammalian gut source. BMC Microbiol. 2015;15: 1–20.

31. Sheng S, Yan S, Chen J, Zhang Y, Wang Y, Qin Q, et al. Gut microbiome is associated with metabolic syndrome accompanied by elevated gamma-glutamyl transpeptidase in men. Front Cell Infect Microbiol. 2022;12. doi:10.3389/fcimb.2022.946757

32. Sedghi L, DiMassa V, Harrington A, Lynch SV, Kapila YL. The oral microbiome: Role of key organisms and complex networks in oral health and disease. Periodontol 2000. 2021;87: 107–131.

33. Ahmed U, Chatterjee T, Kandula M. Polyarteritis Nodosa: an unusual case of paraneoplastic process in renal cell carcinoma. Journal of Community Hospital Internal Medicine Perspectives. 2020;10: 73.

34. De Sordi L, Khanna V, Debarbieux L. The Gut Microbiota Facilitates Drifts in the Genetic Diversity and Infectivity of Bacterial Viruses. Cell Host Microbe. 2017;22: 801–808.e3.

35. Carr VR, Shkoporov A, Hill C, Mullany P, Moyes DL. Probing the Mobilome: Discoveries in the Dynamic Microbiome. Trends Microbiol. 2021;29: 158–170.

36. Dutilh BE, Cassman N, McNair K, Sanchez SE, Silva GGZ, Boling L, et al. A highly abundant bacteriophage discovered in the unknown sequences of human faecal metagenomes. Nat Commun. 2014;5: 1–11.

37. Kolde R, Franzosa EA, Rahnavard G, Hall AB, Vlamakis H, Stevens C, et al. Host genetic variation and its microbiome interactions within the Human Microbiome Project. Genome Med. 2018;10: 6.

38. Luca F, Kupfer SS, Knights D, Khoruts A, Blekhman R. Functional Genomics of Host-Microbiome Interactions in Humans. Trends Genet. 2018;34: 30.

39. Zhang Y, Wang X, Li H, Ni C, Du Z, Yan F. Human oral microbiota and its modulation for oral health. Biomed Pharmacother. 2018;99: 883–893.

40. Le Bars P, Matamoros S, Montassier E, Le Vacon F, Potel G, Soueidan A, et al. The oral cavity microbiota: between health, oral disease, and cancers of the aerodigestive tract. Can J Microbiol. 2017;63: 475–492.

41. Belstrøm D. The salivary microbiota in health and disease. J Oral Microbiol. 2020;12: 1723975.

42. Willis JR, Gabaldón T. The Human Oral Microbiome in Health and Disease: From Sequences to Ecosystems. Microorganisms. 2020;8. doi:10.3390/microorganisms8020308

43. Matsue M, Mori Y, Nagase S, Sugiyama Y, Hirano R, Ogai K, et al. Measuring the Antimicrobial Activity of Lauric Acid against Various Bacteria in Human Gut Microbiota Using a New Method. Cell Transplant. 2019;28: 1528–1541.

44. Vernocchi P, Del Chierico F, Putignani L. Gut Microbiota Profiling: Metabolomics Based Approach to Unravel Compounds Affecting Human Health. Front Microbiol. 2016;7: 1144.

45. Yang J, Pu J, Lu S, Bai X, Wu Y, Jin D, et al. Species-Level Analysis of Human Gut Microbiota With Metataxonomics. Front Microbiol. 2020;11: 2029.

46. Jumas-Bilak E, Carlier JP, Jean-Pierre H, Mory F, Teyssier C, Gay B, et al. Acidaminococcus intestini sp. nov., isolated from human clinical samples. Int J Syst Evol Microbiol. 2007;57. doi:10.1099/ijs.0.64883-0

47. Loubinoux J, Bronowicki J-P, Pereira IAC, Mougenel J-L, Faou AE. Sulfate-reducing bacteria in human feces and their association with inflammatory bowel diseases. FEMS Microbiol Ecol. 2002;40: 107–112.

48. Lu J, Nogi Y, Takami H. Oceanobacillus iheyensis gen. nov., sp. nov., a deep-sea extremely halotolerant and alkaliphilic species isolated from a depth of 1050 m on the Iheya Ridge. FEMS Microbiol Lett. 2001;205: 291–297.

49. Froidurot A, Julliand V. Cellulolytic bacteria in the large intestine of mammals. Gut Microbes. 2022;14: 2031694.

50. Coutinho TA, Venter SN. Pantoea ananatis: an unconventional plant pathogen. Mol Plant Pathol. 2009;10: 325–335.

51. Arashima Y, Kumasaka K, Okuyama K, Kawabata M, Tsuchiya T, Kawano K, et al. [Clinicobacteriological study of Pasteurella multocida as a zoonosis (1). Condition of dog and cat carriers of Pasteurella, and the influence for human carrier rate by kiss with the pets]. Kansenshogaku Zasshi. 1992;66: 221–224.

52. Dehoux P, Marvaud JC, Abouelleil A, Earl AM, Lambert T, Dauga C. Comparative genomics of Clostridium bolteae and Clostridium clostridioforme reveals species-specific genomic properties and numerous putative antibiotic resistance determinants. BMC Genomics. 2016;17: 819.

53. Li Q, Zhou F, Su Z, Li Y, Li J.: A Confirmed Calcifying Bacterium With a Potentially Important Role in the Supragingival Plaque. Front Microbiol. 2022;13: 940643.

54. Ezeji JC, Sarikonda DK, Hopperton A, Erkkila HL, Cohen DE, Martinez SP, et al. Parabacteroides distasonis: intriguing aerotolerant gut anaerobe with emerging antimicrobial resistance and pathogenic and probiotic roles in human health. Gut Microbes. 2021;13: 1922241.

55. Sarshar M, Behzadi P, Scribano D, Palamara AT, Ambrosi C. Acinetobacter baumannii: An Ancient Commensal with Weapons of a Pathogen. Pathogens. 2021;10. doi:10.3390/pathogens10040387

56. Cassir N, Benamar S, La Scola B. Clostridium butyricum: from beneficial to a new emerging pathogen. Clin Microbiol Infect. 2016;22: 37–45.

57. Krawczyk B, Wityk P, Gałęcka M, Michalik M. The Many Faces of Enterococcus spp.-Commensal, Probiotic and Opportunistic Pathogen. Microorganisms. 2021;9. doi:10.3390/microorganisms9091900

58. Parker BJ, Wearsch PA, Veloo ACM, Rodriguez-Palacios A. The Genus Alistipes: Gut Bacteria With Emerging Implications to Inflammation, Cancer, and Mental Health. Front Immunol. 2020;11: 906.

59. Mingeot-Leclercq MP, Glupczynski Y, Tulkens PM. Aminoglycosides: activity and resistance. Antimicrob Agents Chemother. 1999;43: 727–737.

60. Shakil S, Khan R, Zarrilli R, Khan AU. Aminoglycosides versus bacteria--a description of the action, resistance mechanism, and nosocomial battleground. J Biomed Sci. 2008;15: 5–14.

61. Reeves AZ, Campbell PJ, Sultana R, Malik S, Murray M, Plikaytis BB, et al. Aminoglycoside cross-resistance in Mycobacterium tuberculosis due to mutations in the 5’ untranslated region of whiB7. Antimicrob Agents Chemother. 2013;57: 1857–1865.

62. Hormeño L, Ugarte-Ruiz M, Palomo G, Borge C, Florez-Cuadrado D, Vadillo S, et al. Genes Encoding Aminoglycoside O-Nucleotidyltransferases Are Widely Spread Among Streptomycin Resistant Strains of and. Front Microbiol. 2018;9: 2515.

63. Hooper DC. Emerging mechanisms of fluoroquinolone resistance. Emerg Infect Dis. 2001;7: 337–341.

64. Robicsek A, Jacoby GA, Hooper DC. The worldwide emergence of plasmid-mediated quinolone resistance. Lancet Infect Dis. 2006;6: 629–640.

65. Cheng AFB, Yew WW, Chan EWC, Chin ML, Hui MMM, Chan RCY. Multiplex PCR amplimer conformation analysis for rapid detection of gyrA mutations in fluoroquinolone-resistant Mycobacterium tuberculosis clinical isolates. Antimicrob Agents Chemother. 2004;48: 596–601.

66. Schweizer I, Peters K, Stahlmann C, Hakenbeck R, Denapaite D. Penicillin-binding protein 2x of Streptococcus pneumoniae: the mutation Ala707Asp within the C-terminal PASTA2 domain leads to destabilization. Microb Drug Resist. 2014;20: 250–257.

67. Arnold BJ, Huang I-T, Hanage WP. Horizontal gene transfer and adaptive evolution in bacteria. Nat Rev Microbiol. 2021;20: 206–218.

68. Sparks IL, Derbyshire KM, Jacobs WR Jr, Morita YS. Mycobacterium smegmatis: The Vanguard of Mycobacterial Research. J Bacteriol. 2023;205: e0033722.

69. Mock F, Kretschmer F, Kriese A, Böcker S, Marz M. Taxonomic classification of DNA sequences beyond sequence similarity using deep neural networks. Proc Natl Acad Sci U S A. 2022;119: e2122636119.

70. Peters SL, Borges AL, Giannone RJ, Morowitz MJ, Banfield JF, Hettich RL. Experimental validation that human microbiome phages use alternative genetic coding. Nat Commun. 2022;13: 5710.

71. Hammerling MJ, Ellefson JW, Boutz DR, Marcotte EM, Ellington AD, Barrick JE. Bacteriophages use an expanded genetic code on evolutionary paths to higher fitness. Nat Chem Biol. 2014;10: 178–180.

72. Shim H, Shivram H, Lei S, Doudna JA, Banfield JF. Diverse ATPase Proteins in Mobilomes Constitute a Large Potential Sink for Prokaryotic Host ATP. Front Microbiol. 2021;12: 691847.

73. Park H-M, Park Y, Vankerschaver J, Van Messem A, De Neve W, Shim H. Rethinking Protein Drug Design with Highly Accurate Structure Prediction of Anti-CRISPR Proteins. Pharmaceuticals. 2022;15: 310.

74. Shim H. Investigating the genomic background of CRISPR-Cas genomes for CRISPR-based antimicrobials. arXiv [q-bio.GN]. 2022. Available: http://arxiv.org/abs/2202.07171

75. Park H-M, Park Y, Berani U, Bang E, Vankerschaver J, Van Messem A, et al. In silico optimization of RNA-protein interactions for CRISPR-Cas13-based antimicrobials. Biol Direct. 2022;17: 27.

76. Kasianowicz JJ, Brandin E, Branton D, Deamer DW. Characterization of individual polynucleotide molecules using a membrane channel. Proc Natl Acad Sci U S A. 1996;93: 13770–13773.

77. Grädel C, Terrazos Miani MA, Barbani MT, Leib SL, Suter-Riniker F, Ramette A. Rapid and Cost-Efficient Enterovirus Genotyping from Clinical Samples Using Flongle Flow Cells. Genes. 2019;10. doi:10.3390/genes10090659

78. Antimicrobial Resistance Collaborators. Global burden of bacterial antimicrobial resistance in 2019: a systematic analysis. Lancet. 2022;399: 629–655.

79. Nicholls SM, Quick JC, Tang S, Loman NJ. Ultra-deep, long-read nanopore sequencing of mock microbial community standards. Gigascience. 2019;8: giz043.

80. Street TL, Barker L, Sanderson ND, Kavanagh J, Hoosdally S, Cole K, et al. Optimizing DNA Extraction Methods for Nanopore Sequencing of Neisseria gonorrhoeae Directly from Urine Samples. J Clin Microbiol. 2020;58. doi:10.1128/JCM.01822-19

81. Marquet M, Zöllkau J, Pastuschek J, Viehweger A, Schleußner E, Makarewicz O, et al. Evaluation of microbiome enrichment and host DNA depletion in human vaginal samples using Oxford Nanopore’s adaptive sequencing. Sci Rep. 2022;12: 1–10.

82. Martin S, Heavens D, Lan Y, Horsfield S, Clark MD, Leggett RM. Nanopore adaptive sampling: a tool for enrichment of low abundance species in metagenomic samples. Genome Biol. 2022;23: 1–27.

83. Hingamp P, Grimsley N, Acinas SG, Clerissi C, Subirana L, Poulain J, et al. Exploring nucleo-cytoplasmic large DNA viruses in Tara Oceans microbial metagenomes. ISME J. 2013;7: 1678–1695.

84. Shim H. Feature Learning of Virus Genome Evolution With the Nucleotide Skip-Gram Neural Network. Evol Bioinform Online. 2019;15: 1176934318821072.

